# Strategies of resource sharing in clonal plants: A conceptual model and an example of contrasting strategies in two closely related species

**DOI:** 10.1101/2024.04.02.587797

**Authors:** Jana Duchoslavová, Jan Jansa

**Affiliations:** Department of Botany, Faculty of Science, Charles University, Benátská 2, 128 43 Praha 2, Czech Republic; Institute of Microbiology, Czech Academy of Sciences, Vídeňská 1083, 142 20 Praha 4, Czech Republic

**Keywords:** Carbon, clonal plants, development, light, nitrogen, physiological integration, stable isotopes, translocation

## Abstract

**Background and Aims:** Clonal growth helps plants to cope with environmental heterogeneity through resource integration via connecting organs. Such integration is considered to balance heterogeneity by the translocation of resources from rich to poor patches. However, such an ‘equalisation’ strategy is only one of several possible strategies, as we discuss in a brief conceptual analysis. Under certain conditions, a strategy emphasising acropetal movement and exploration of new areas or a strategy accumulating resources in older ramets may be preferred. The optimal strategy may be determined by environmental conditions, such as resource availability and level of light competition. Therefore, species from different habitats may exhibit distinct resource translocation strategies.

**Methods:** Resource translocation was compared between two closely related species from different habitats with contrasting productivity. The study examined the bidirectional translocation of carbon and nitrogen in pairs of mother and daughter ramets grown under light heterogeneity (one ramet shaded) at two developmental stages using stable-isotope labelling.

**Key results:** At the early developmental stage, both species translocated resources toward daughters and the translocation was modified by shading. Later, the species of low-productivity habitats, *Fragaria viridis*, translocated carbon to shaded ramets, according to the ‘equalisation’ strategy. The species of high-productivity habitats, *Potentilla reptans*, did not support shaded older ramets. Nitrogen translocation remained mainly acropetal in both species. These findings confirmed our expectations.

**Conclusions:** The two studied species exhibited different translocation strategies, which may be linked to the habitat conditions experienced by each species. The ‘equalisation’ strategy may occur in habitats with lower productivity and lower light competition, than the strategy emphasising acropetal movement. The results indicate that we need to consider different possible strategies. We emphasise the importance of bidirectional tracing in translocation studies and the need for further studies to investigate the translocation patterns in species from contrasting habitats.

## Introduction

As sessile organisms, plants cope with environmental heterogeneity, e.g. in resource availability, at a very local scale. Plant species have evolved various traits, including specific morphological adaptations or phenotypic plasticity, to survive resource limitations. One of these strategies is clonal growth, which allows plants to spread horizontally via clonal organs such as stolons and rhizomes. Such growth enables plants to explore new adjacent patches and integrate resources from a larger area through the interconnected plant body, thereby adjusting to or benefiting from environmental heterogeneity (Marshall, 1990; Song *et al*., 2013).

Resource sharing among interconnected ramets (i.e. potentially independent plant parts rooting at different points) is considered one of the main advantages of clonal growth. Clonal organs allow ramets to share water, mineral nutrients, and photosynthates and to transport signal molecules from ramet initialisation until the connection ends (Pitelka & Ashmun, 1985; Marshall, 1990; Alpert *et al*., 2002). Initially, newly developing daughter ramets are supported by mother ramets because their resource demands are not covered by their limited resource uptake capacity. This support is analogous to maternal provision to seeds in sexual reproduction (Hartnett & Bazzaz, 1983; Bullock *et al*., 1994; Wijesinghe, 1994). The initial maternal support, which appears to be universal across species, may change later in ramet ontogeny depending on a species’s resource-sharing strategy and resource availability in the environment (Pitelka & Ashmun, 1985; Duchoslavová & Jansa, 2018; Ma *et al*., 2021). Whereas persisting resource translocation among developed ramets may not be beneficial in stable, homogeneous habitats, it may be advantageous when resources are distributed heterogeneously in space or time (e.g., Evans, 1991; Alpert 1999; Wang *et al*., 2021).

However, predicting translocation patterns under environmental heterogeneity is not straightforward, as different magnitudes and directions of resource translocation may be optimal in different contexts (Pitelka & Ashmun, 1985; Alpert, 1999). Correspondingly, both intra- and interspecific variation in resource translocation have been shown in clonal species (Alpert, 1999; Xu *et al*., 2010). In this paper, we outline several possible distinct resource translocation strategies (known or expected) among developed ramets in heterogeneous environments, and we show existence of two different resource-sharing patterns in two closely related stoloniferous species. We propose names for the resource-sharing strategies, as we are not aware of any existing terminology that would be appropriate for this purpose, and we feel that the potentially contrasting patterns of resource translocation deserve to be named.

### Hypothetical types of resource sharing

In the first possible strategy, developed ramets growing in resource-poor patches may be supported by ramets growing in resource-rich patches, regardless of their ontogenetic position (i.e. both acropetally and basipetally; Fig. 1a). We call this the Equalisation strategy of resource sharing. It has been demonstrated in many studies (e.g., Alpert & Mooney, 1986; Evans, 1991; Shumway, 1995), and it has been often implicitly considered to be synonymous with clonal integration in the recent literature (e.g., Liu *et al.,* 2016, Wang *et al*., 2021). However, simple support of resource-limited ramets may not be the optimal resource-sharing strategy in all conditions, and other patterns of resource sharing may occur (Pitelka & Ashmun, 1985; Alpert *et al*., 2002). In a successful resource-sharing strategy, translocating a certain amount of a resource should produce more fitness benefits for the recipient ramets than costs for donor ramets, resulting in higher overall fitness of the whole clonal fragment. This strongly depends on the shape of the resource–fitness relationship, which may vary under different conditions (Eriksson & Jerling, 1990; Alpert, 1999). For example, if resource-poor patches are too unfavourable or if the chance of future improvements in resource availability is too low, relative gain of the receiving ramet may not outweigh the cost of translocation.

**Figure 1.**
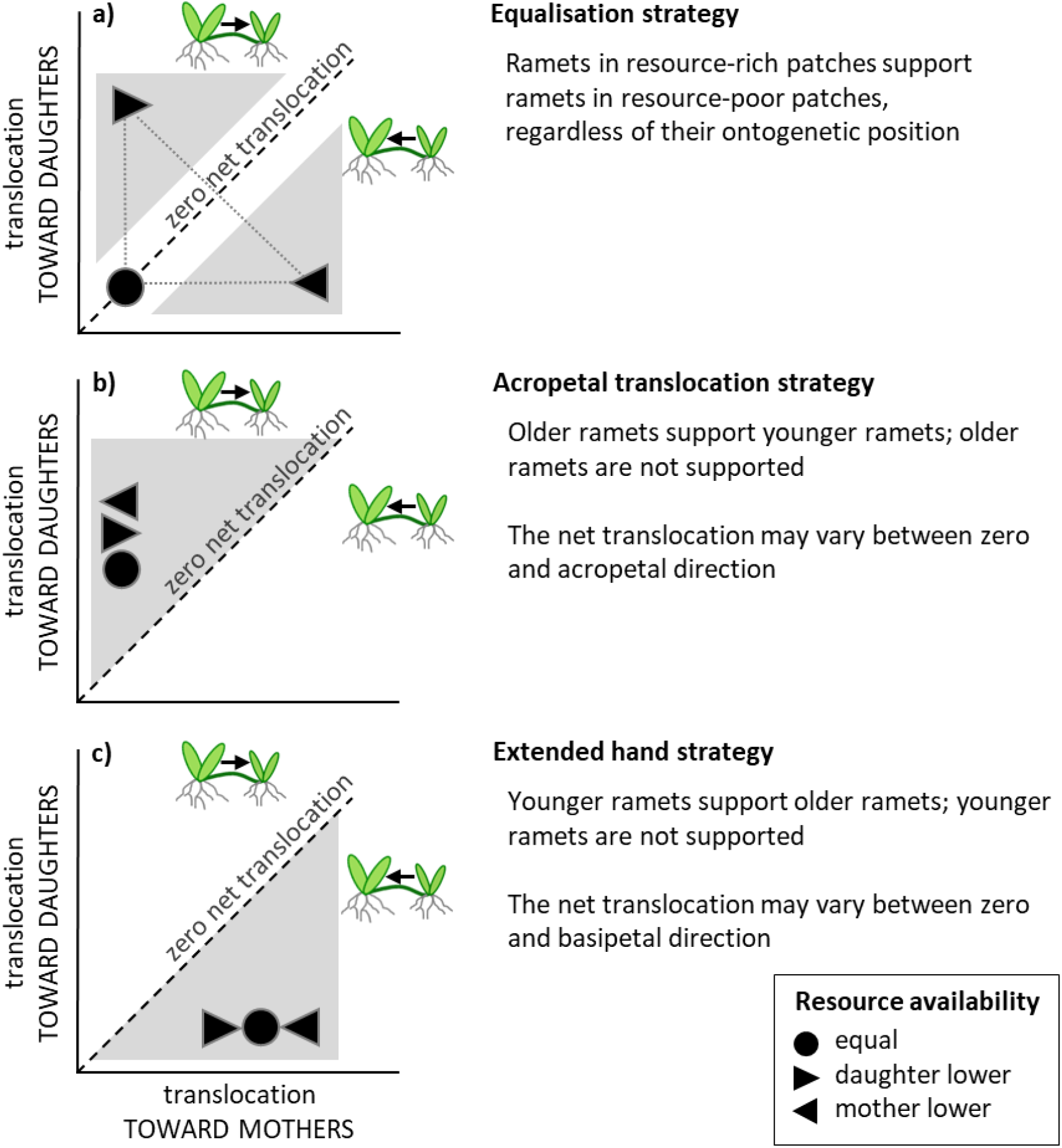
Hypothetical **two-way translocation plots** for the proposed resource-sharing strategies between mother and daughter ramets in the later developmental stage. The dashed line indicates zero net translocation. Translocation toward daughters prevails in the zone above the dashed line, while translocation toward mothers prevails in the zone below the dashed line. Grey triangles indicate zones in which the values may range.

Alternatively, plants may employ a resource-sharing strategy based on the acropetal horizontal movement of the plant body and exploration of new areas rather than prolonged support of resource-limited ramets. This strategy may be preferable especially when the environment is patchy also in time (Pitelka & Ashmun, 1985; Gardner & Mangel, 1999), and when resources may get locally depleted. We call this the Acropetal translocation strategy, and it is characterized by mainly acropetal translocation of resources – developmentally older ramets are not supported even if they are resource-limited (Noble & Marshall, 1983; Xiao *et al*., 2011; Fig. 1b). Support of younger ramets growing at lower resource level and no similar support of older ramets was observed in several species (Slade & Hutchings, 1987; van Kleunen & Stuefer, 1999; Portela *et al*., 2021). Such translocation pattern may promote plant invasion into vegetated areas (Xiao *et al*., 2011) or clonal spread through unfavourable areas to colonize potential new favourable patches. However, acropetal translocation may not only equalise resource availability. Younger ramets growing at higher resource level might be supported preferentially to maximise profit from favourable patches. Indications of such translocation pattern have been described, for example, in *Fragaria chiloensis*, where offspring ramets in nutrient-rich patches benefited from clonal integration with mothers (Alpert, 1996), or in *Agrostis stolonifera*, where carbon tended to be translocated acropetally from shaded to unshaded ramets (Duchoslavová & Jansa, 2018). Mainly acropetal direction of resource translocation may be driven by physiological constraints in some species (Stuefer, 1996), although the possibility of basipetal translocation has been demonstrated repeatedly (Tietema & van der Aa, 1981; Shumway, 1995; Jonsdottir & Watson, 1997). Resource sharing strategies may be coupled with morphological and behavioural characteristics of plants. For example, the structural blue-print that determines constant mobility of clonal species may be associated with the Acropetal translocation strategy (*Glechoma hederacea*; Huber *et al.,* 1999). Species with the Acropetal translocation strategy may have lower persistence of ramets compared to species with the Equalisation strategy. In addition, they may exhibit stronger changes in morphology of clonal growth to escape from unfavourable conditions, active foraging for available resources and horizontal spread toward favourable patches (Price & Hutchings, 1996).

Moreover, daughter ramets could hypothetically be used as an extended hand of a mother ramet for a period of time, supporting the mother ramet’s resource demands (Extended Hand strategy; Guo et al., 2020, Pinno & Wilson, 2014; Fig. 1c). Such a strategy may be preferable when concentrating resources in the mother ramet brings benefits for the entire clonal fragment, such as when the mother ramet is flowering while the daughter ramets remain vegetative (Guo et al., 2020). Presumably, this strategy may occur in species with long persisting ramets and young ramets remaining rather small and vegetative in the first year of their development. Due to internal sources of resource availability gradients, such translocation may be rather independent of environmental resource heterogeneity. We expect this strategy to be particularly effective for exploration of soil-borne resources which might get depleted by older ramets.

Finally, plants might exhibit no net translocation between developed ramets (Zero net translocation strategy). In this case, the clonal connection may be interrupted, or it may persist only as a remnant of the early translocation. Alternatively, the connection can be actively maintained in case of need, such as high stress, regeneration after disturbance.

Which resource-sharing strategy is preferable may be determined by the nature of resource distribution and of plant competition in the environment (Pitelka & Ashmun, 1985). In terms of light heterogeneity, this can have different consequences depending on the context. Light competition is size asymmetric, which means that the tallest plants gain disproportionately more light. Therefore, in habitats where the surrounding vegetation is typically higher than the focal plant, shaded ramets have a low chance of reaching the top of the canopy, and it may not be advantageous to invest resources to maintain them. Light is thus scarce and patchily distributed in vegetation gaps for such shorter plants. Accordingly, the Acropetal translocation strategy may be preferred if there is major horizontal heterogeneity in light availability, as it might enable rapid spread to new, potentially unshaded patches. On the other hand, when all individuals have a chance to reach the top of the canopy, as in the case of tall plants or in nutrient-poor habitats with low surrounding vegetation, light is not as scarce and patchy. In such cases, the Equalisation strategy may be preferred since it enables the maintenance of established ramets (as suggested also by Wang *et al*., 2021).

Different resource types have different mechanisms of uptake, translocation and use in plants (e.g. Marshall, 1990), but are still not independent of each other. Translocation of photosynthates and part of nutrients by the phloem is regulated primarily by activity of sinks and sources (Marshall, 1990), and it may be further modified by hormonal control (Alpert *et al*., 2002). In contrast, water and most nutrients are translocated by the transpiration flux in the xylem, which is mainly determined by water potential gradients. Accordingly, translocation of nitrogen between ramets has been shown to depend on transpiration flux together with the nitrogen availability gradient (de Kroon et al. 1998). However, plants respond to resource limitation by adjusting their biomass allocation, morphology, physiology, and architecture, which affects the uptake and presumably translocation of several resources (Freschet et al., 2018). Moreover, translocation of photosynthates from unshaded to shaded conditions may be accompanied by reciprocal translocation of water due to higher evapotranspiration under unshaded conditions (Stuefer et al., 1994). Thus, the uptake and translocation of different types of resources in clonal plants are clearly not independent, and examination of multiple resource translocation could help to reveal integrated plant responses to resource limitation.

It is not yet clear how often different strategies of resource translocation occur in clonal plants and how they depend on environmental conditions. While experiments evaluating the effects of clonal integration on ramet biomass provide a valuable indication of translocation patterns, they usually do not separate the effects of translocation at early and late developmental stages, which may be completely reversed (but see Xu *et al*., 2012; Ma *et al*., 2021). In our opinion, the most appropriate way to directly evaluate resource-sharing patterns among established ramets is to trace labelled resources in both directions, which reveals the translocation pattern at a given stage. Nevertheless, this approach is rarely used (but see Duchoslavová & Jansa, 2018, Zhai et al., 2022).

### Experimental approach

We chose two closely related stoloniferous species from the Rosaceae family and compared their resource-sharing strategies under light heterogeneity. We chose the species because they exhibit contrasting habitat preferences. *Fragaria viridis* inhabits dry grasslands with low vegetation, whereas *Potentilla reptans* can grow in more fertile habitats with higher vegetation, such as mesic meadows. Therefore, *F. viridis* is likely exposed to higher levels of light and lower availability of belowground resources in its natural habitats than *P. reptans* and may exhibit a different resource-sharing strategy.

In this study, we grew pairs of mother and daughter ramets of these species with one ramet unshaded and one ramet shaded by green shade simulating light competition. We traced labelled carbon and nitrogen in both directions to determine the resource-sharing strategies of the species. Whereas carbon translocation is directly linked to light heterogeneity, nitrogen tracing helped us gain a more complex picture of plants’ resource economy, nutrient uptake capacity and relative resource limitation. In addition, we examined resource-sharing in early and later developmental stages of daughter ramets (2 weeks and 8 weeks after daughter-ramet initiation, respectively) to observe the change in resource-sharing patterns during ramet development.

We expected both species to translocate carbon and nitrogen toward daughter ramets in the early developmental stage. We hypothesized that unshaded daughters would form stronger sinks for nitrogen and therefore import more nitrogen than shaded daughters. In the later developmental stage, we expected carbon translocation to switch to the Equalisation strategy in *F. viridis* and to the Acropetal strategy in *P. reptans*. We hypothesized that nitrogen translocation would be directed toward unshaded mother or daughter ramets which are not limited by carbon, and likely form stronger sinks for nitrogen.

## Materials and Methods

### Species

Both experimental species belong to the Rosaceae family. They are perennial, form rosettes of leaves and spread horizontally via stolons with similar lateral spreading distances (20 cm for *F. viridis* and 18 cm for *P. reptans*; Klimešová *et al*., 2017). Connections among ramets persist for the whole vegetative season. *Fragaria viridis* is common in pastures, dry grasslands, rocky steppes, and open forest edges (Slavík *et al*., 1995; Ellenberg’s indicator values of 7 for light and 4 for nitrogen; Chytrý *et al*., 2018). *Potentilla reptans* occurs in mesic meadows, stream banks, ruderal areas, fields and forest edges (Slavík *et al*., 1995; Ellenberg indicator values of 6 for light and 6 for nitrogen; Chytrý *et al*., 2018). Therefore, *P. reptans* inhabits more nutrient-rich habitats than *F. viridis* and tolerates more shading.

Three genotypes per species were collected in the field from localities around Prague (Czech Republic) in 2016 and grown as source plants in an experimental garden at Charles University in Prague. Different genotypes were collected at locations at least 100 m apart from each other.

### Initial cultivation

The stolons of source plants were placed on wet perlite to initiate rooting in late May 2017. After 2 weeks, individual ramets were separated and planted in 2 L pots with a mixture of washed sand and zeolite (1:1). The substrate was fertilised with slow-release fertiliser (Substral Osmocote for garden, 7 g per pot). These ramets are referred to below as the mother ramets. The pots were placed in a greenhouse equipped with supplemental lighting from 400 W metal halide lamps providing a minimum of 200 μmol m^−2^ s^−1^ photosynthetically active radiation (PAR), extending the daylight period to 14 h. They were watered two times a day with tap water. Dead ramets were replaced by new ones until mid-June 2017. The plants were treated with insecticide to protect them against pests (specifically, *Tetranychus urticae*; the insecticides Nissorun and Karate were used as per the manufacturers’ recommendations, www.agrobio.cz).

By early July, the mother ramets had produced one or more new stolons bearing several new rosettes of leaves. The initial size of the mother ramets was measured 1 day before the initialisation of daughter-ramet rooting as the number of leaves and number of stolons. The rooting of daughter ramets was initiated between 13–16 July 2017 by placing the longest stolon of each mother ramet in an adjacent vacant pot. The tips of these stolons were kept intact, and all the other stolons were left on plants, but they were not allowed to root (similarly to Alpert, 1999). All the unrooted stolons were kept in the same shade treatment as their associated ramet. Therefore, by a mother ramet we mean a developmentally older rooted rosette of leaves and all the unrooted stolons associated with this rosette, and by a daughter ramet we mean a developmentally younger rooted rosette of leaves and all the associated unrooted stolons. For logistic reasons, the initialisation of daughter-ramet rooting, as well as labelling and harvesting, were conducted over 4 subsequent days. This made it possible to maintain the same intervals between the initialisation of daughter-ramet rooting and harvesting for all plants. Therefore, the ramet pairs processed at each harvest step (i.e. for a given developmental stage) were divided into 4 time-blocks with 1-d differences in rooting initialisation/harvest. The treatments were represented evenly among the blocks.

### Experimental design

We used an experimental design with three shading treatments: (i) both ramets in full light, (ii) daughters shaded, and (iii) mothers shaded (Fig. 2). A shading cloth was installed 1 day before the initialisation of daughter-ramet rooting. We used the shading cloth combined with 3-cm-wide strips of green foil (LeeFilters Fern Green 122, www.leeflters.com) to shade the plants from the top and all sides, thus simulating the changes in both light quantity and quality caused by aboveground competition (80% PAR reduction and 30% reduction of the red to far-red ratio, as measured by the spectroradiometer Avaspec-2048, VA300; see Fig. S1 for a picture of the shading setup; Duchoslavová & Jansa, 2018).

**Figure 2.**
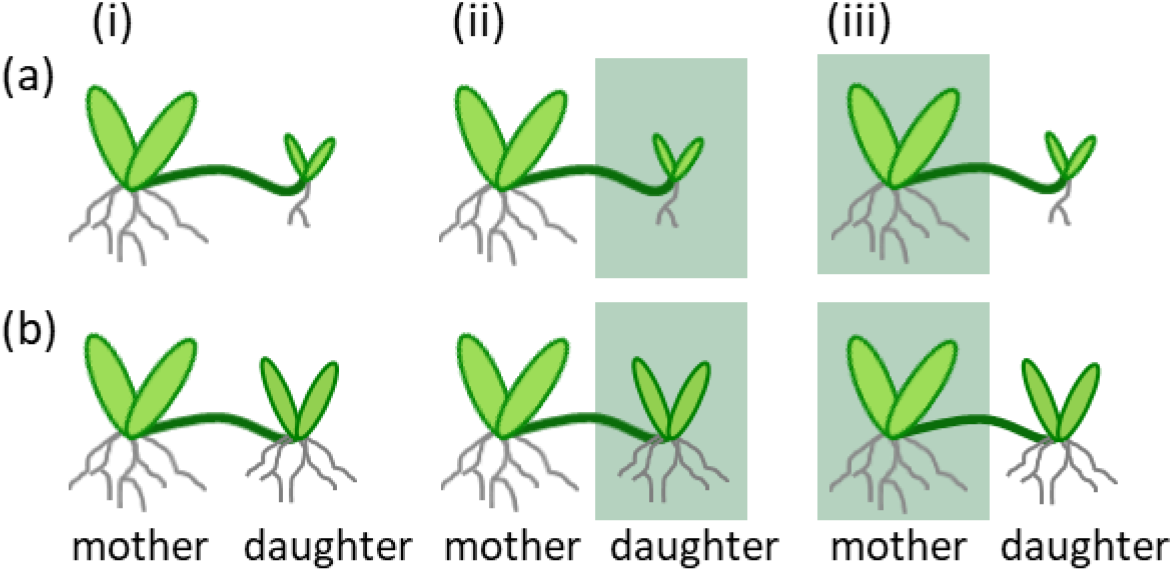
Three shading treatments with green shade were used in the experiment: full light (i), daughter shaded (ii) and mother shaded (iii). The plants were pulse-labelled by stable isotopes of C and N and harvested in two developmental stages of daughter ramets: (a) the early developmental stage (2 weeks after initialisation of daughter rooting) and (b) the later developmental stage (8 weeks after initialisation of daughter rooting). Note that new unrooted stolons forming on both mother and daughter ramets are not depicted in this scheme.

The bidirectional translocation of carbon and nitrogen between mothers and daughters was examined by stable-isotope pulse labelling (using ^13^C and ^15^N) at two developmental stages of the daughters: 2 weeks after the initialisation of rooting (early developmental stage) and 8 weeks after the initialisation of rooting (later developmental stage). In each ramet pair, either translocation to daughter ramets or translocation to mother ramets was measured. Four replicates were used for each combination of species, shading treatment, measured direction of translocation and time of labelling.

### Stable-isotope labelling

Plants were pulse-labelled simultaneously by nitrogen (^15^N) and carbon (^13^C) according to a protocol used in a previous experiment (Duchoslavová & Jansa, 2018). Labelling began at approximately 8:00 a.m. Nitrogen was applied directly to the substrate of a labelled ramet by a syringe in the form of doubly labelled ammonium nitrate (99 atom% ^15^N, 15 mg per pot). Carbon was applied in the form of ^13^CO_2_. Pots with labelled ramets were enclosed in plexiglass chambers equipped with a fan to mix the inner atmosphere, and 20 ml of phosphoric acid (20%, w:v) was injected into a vial with ^13^C-enriched sodium carbonate (99% atom% ^13^C, 0.3 g per pot) to release ^13^CO_2._ The calculated initial ^13^CO_2_ concentration inside the chambers reached 3,100 ppm. The ramets were allowed to assimilate labelled carbon for 2 hours, and the remaining CO_2_ in the atmosphere was then scrubbed by circulating the air through an NaOH solution (0.1 M, 200 ml). The labelling took place in full sunlight supplemented by additional light from diode lamps (LumiGrow Pro Series Pro 325 LED Lighting Systems, photosynthetic photon flux density 333 μmol m-^2^ s^-1^ at a distance of 1 m), and the plants were returned to experimental shading conditions immediately after the labelling period.

Labelled ramet pairs were harvested exactly 2 days after labelling. This period enables labelled elements to go through the entire day cycle but still reflects actual (short-term) rather than cumulative (long-term) resource translocation. Mother and daughter ramets were separated, and the roots were washed and separated from the shoots. Unrooted stolons were kept on the plants and considered as part of the shoots. The plant material was then dried (65 °C for 2–3 days), weighed and ground to a fine powder using a ball mill (MM200, Retsch, Haan, Germany) before the elemental and isotopic analyses. The N and C concentrations and isotopic composition of the two elements were measured using an elemental analyser (Flash EA 2000) coupled with an isotope ratio mass spectrometer (Delta V Advantage, ThermoFisher Scientific, Waltham, MA, USA).

To calculate the amount of ^13^C and ^15^N originating from labelling pulse (i.e. excess ^13^C or ^15^N), F-ratios, i.e. the proportion of the heavy isotope in the total amount of the element, were calculated as *R_S_*/(*R_S_* +1), where *R_S_* is the molar isotope ratio in a sample (^15^N/^14^N or ^13^C/^12^C). The amount of total carbon or nitrogen a sample (*C*, in moles) was then calculated as follows:

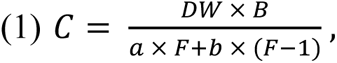

where *DW* is the dry weight of the sample, *B* is the molar concentration of carbon or nitrogen in a sample, *a* is 12 for carbon and 14 for nitrogen, *b* is 13 for carbon and 15 for nitrogen, and *F* stands for the F-ratio of the respective element in a sample, as specified above. F-ratios of ^13^C or ^15^N originating from pulse-labelling (*F_D_*) were calculated as

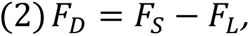

where *F_S_* is a sample’s F-ratio and *F_L_* is a limit F-ratio below which we cannot detect the stable-isotope enrichment with a sufficient confidence. This value was calculated from the F-ratios of the unlabelled control samples as 99th percentile of an approximated normal distribution. Therefore, F-ratio values above this limit would come from this distribution with probability less than 1 %. Negative F_D_ values were replaced by zeroes.

The amount of ^13^C and ^15^N originating from pulse-labelling (*E*, in moles) was finally calculated as

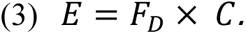

The amount of ^13^C and ^15^N originating from pulse-labelling in unlabelled ramets (roots and shoots combined) is further referred to as the amount of translocated ^13^C and ^15^N.

As all ramets were labelled in full-light conditions to allow for sufficient incorporation of ^13^C into plant biomass, excess ^13^C values of shaded ramets and corresponding amounts of translocated ^13^C might be overestimated. However, the qualitative pattern of ^13^C translocation over the subsequent two-day period and fractions of ^13^C exported to the other ramet were presumably unaffected by the labelling conditions.

### Data analyses

We performed statistical analyses in the R statistical environment using linear mixed-effects models (lmerTest package; Kuznetsova *et al*., 2017). These models enabled the description of the data’s hierarchical structure, which was caused by the use of several genotypes. P-values were provided via Satterthwaite’s degrees of freedom method. For the analysis of ramet biomass, the effects of initial number of leaves, initial number of stolons, ramet (mother/ daughter), shading, species, and their interactions were modelled in a separate model for each developmental stage. Genotype identity was included as a random effect affecting the intercept. The biomass was log-transformed in order to meet the model assumptions. For the analyses of translocation, the effects of species, traced translocation direction, shading and their interactions on translocated ^13^C and ^15^N for the two developmental stages were modelled in four separate models (one model for each combination of resource and developmental stage). Genotype identity was included as a random effect affecting the intercept, and its effect was allowed to change with direction and shading. The translocated ^13^C and ^15^N were log-transformed in all models in order to meet the model assumptions.

## Results

### Ramet size and uptake of carbon and nitrogen

At the early developmental stage (i.e. 2 weeks after daughter rooting initialisation), there were no pronounced differences in ramet biomass between the two species. Mothers in the full-light treatment reached 4.6 times higher average dry biomass than daughters. Shading the daughters reduced biomass of both ramets by 40%. Shading the mothers reduced biomass of mothers by 50% in *F. viridis* and by 30% in *P. reptans* while it reduced biomass of daughters only in *F. viridis* by 40% (Fig. 3, Table S1). In terms of carbon uptake, mothers in the full-light treatment assimilated on average 3.1 times more labelled carbon per ramet than daughters in full light. Shading had a similar effect on carbon uptake as it did on biomass (Fig. S2). It should be noted that the carbon uptake values represent the potential uptake capacity of the ramets measured in full light for all ramets. Nitrogen uptake was very low in young daughters, leading to disproportionally higher uptake by mothers (7 and 23.6 times in *F. viridis* and *P. reptans*, respectively). This was likely caused by the markedly lower root mass fraction of daughters in comparison to mothers (Fig. S3). On average, *P. reptans* mothers assimilated 2 times more labelled nitrogen than mothers of *F. viridis* at the time of the first harvest (Fig. S2).

**Figure 3.**
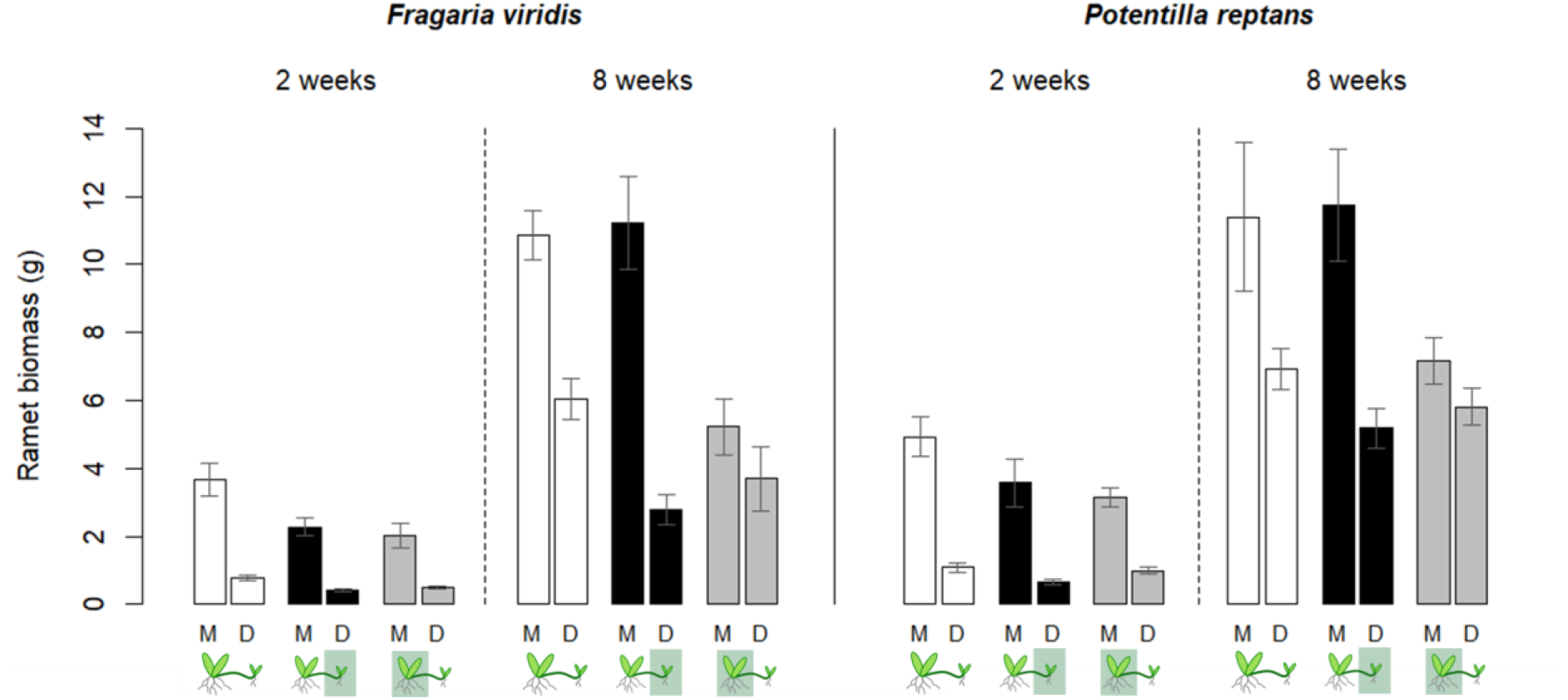
Biomass of mother (M) and daughter (D) ramets in early and later developmental stages of daughter ramets (i.e. 2 and 8 weeks after daughter initialisation). Means and SEM are depicted.

At the later developmental stage (i.e. 8 weeks after daughter rooting initialisation), mothers of *F. viridis* and *P. reptans* in the full-light treatment had 1.9 and 1.5 times higher average dry biomass than daughters, respectively. Shading reduced the biomass of the shaded ramets by 60% times in both ramets of *F. viridis* and by 30% in both ramets of *P. reptans.* Moreover, shading the mothers reduced daughter biomass by 50% in *F. viridis* and by 20% in *P. reptans* (Fig. 3, Table S1). Shading the daughters did not reduce the biomass of mothers in either species (Fig. 3). Despite the differences in biomass, daughter and mother ramets assimilated on average similar amount of labelled carbon in both species. Shading treatments affected the carbon uptake capacity only to a limited extent (Fig. S2). Nitrogen uptake was proportional to root mass of labelled ramets (R^2^ = 0.83, data not shown) and remained higher in mother ramets at the later developmental stage. In the full-light treatment, mothers assimilated on average 1.5 times more labelled nitrogen than daughters in both species, with *P. reptans* assimilating 1.3 times more than *F. viridis*. At the later developmental stage, the effect of shading on nitrogen uptake was similar to its effect on biomass (Figs 3 and S2).

### Carbon translocation

At the early developmental stage, carbon was translocated significantly more from mothers to daughters than from daughters to mothers. This effect was significantly modified by shading treatment. Specifically, shading the mothers increased translocation directionality toward mothers in both species (Table 1, Fig. 4).

**Figure 4.**
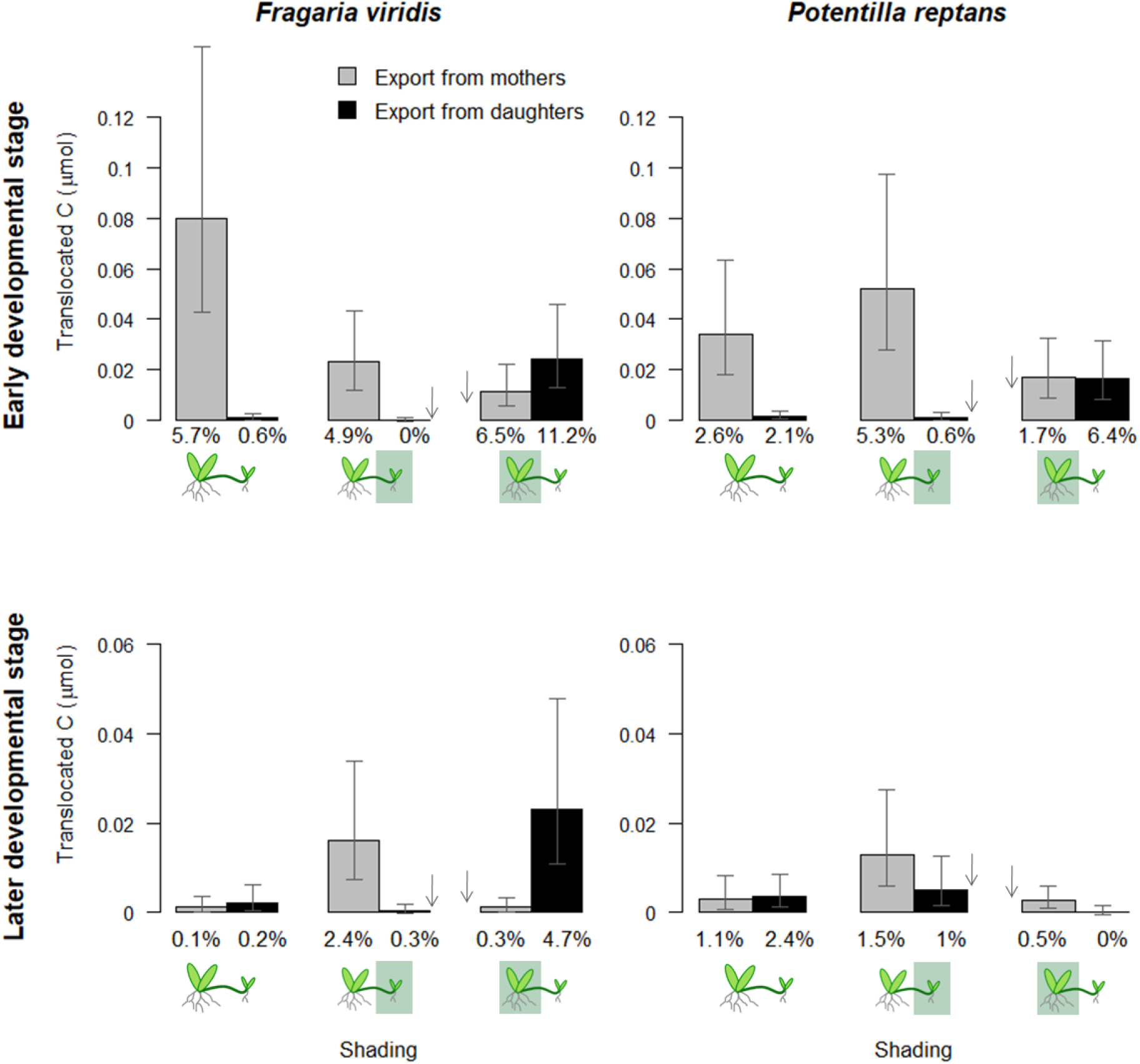
Absolute amounts of translocated ^13^C (means and SEM) from labelled mother to daughter ramets and from labelled daughter to mother ramets. Numbers under bars indicate mean fractions of ^13^C exported from labelled ramets toward unlabelled ramets. Arrows next to the bars indicate which absolute values have possibly been overestimated by the labelling method and the direction where more realistic values may lie. Note that y-axis range for the later developmental stage is half of that for the early developmental stage. Values of absolute amounts are based on linear models with logarithmic transformation (values were back-transformed).

**Table 1.** ANOVA table of the linear mixed-effects models of C and N translocation. Note the corrected values of denominator degrees of freedom (see Methods for details).

| Translocated <sup>13</sup>C |  |  |  |  |  |  |  |  |  |  |  |  |
| --- | --- | --- | --- | --- | --- | --- | --- | --- | --- | --- | --- | --- |
|  |  | First harvest |  |  |  |  |  | Second harvest |  |  |  |  |
|  | d.f. | SumSq | MeanSq | DenDF | Fvalue | Pr(>F) |  | SumSq | MeanSq | DenDF | Fvalue | Pr(>F) |
| Species | 1 | 0.72 | 0.72 | 10.46 | 0.54 | 0.478 |  | 0.03 | 0.03 | 3.49 | 0.02 | 0.907 |
| Direction of translocation | 1 | **48.03** | **48.03** | **31.57** | **35.94** | **<0.001** |  | 0.87 | 0.87 | 29.93 | 0.51 | 0.483 |
| Shading | 2 | 4.09 | 2.05 | 5.28 | 1.53 | 0.299 |  | 3.00 | 1.50 | 8.93 | 0.87 | 0.452 |
| Species x direction | 1 | 0.03 | 0.03 | 31.57 | 0.02 | 0.877 |  | 1.78 | 1.78 | 29.93 | 1.03 | 0.318 |
| Species x shading | 2 | 2.00 | 1.00 | 5.28 | 0.75 | 0.517 |  | 7.34 | 3.67 | 8.93 | 2.12 | 0.176 |
| Direction x shading | 2 | **32.86** | **16.43** | **31.57** | **12.30** | **<0.001** |  | **11.56** | **5.78** | **29.86** | **3.35** | **0.049** |
| Species x direction x shading | 2 | 1.75 | 0.88 | 31.57 | 0.66 | 0.526 |  | **12.95** | **6.47** | **29.86** | **3.75** | **0.035** |
| Translocated <sup>15</sup>N |  |  |  |  |  |  |  |  |  |  |  |  |
|  |  | First harvest |  |  |  |  |  | Second harvest |  |  |  |  |
|  | d.f. | SumSq | MeanSq | DenDF | Fvalue | Pr(>F) |  | SumSq | MeanSq | DenDF | Fvalue | Pr(>F) |
| Species | 1 | **0.86** | **0.86** | **4.57** | **18.22** | **0.010** |  | 3.80 | 3.80 | 5.26 | 2.07 | 0.207 |
| Direction of translocation | 1 | **56.56** | **56.56** | **4.38** | **1200.73** | **<0.001** |  | **42.14** | **42.14** | **30.25** | **22.90** | **<0.001** |
| Shading | 2 | **1.01** | **0.51** | **3.85** | **10.71** | **0.027** |  | 0.20 | 0.10 | 5.70 | 0.05 | 0.948 |
| Species x direction | 1 | **0.63** | **0.63** | **4.38** | **13.26** | **0.019** |  | 4.06 | 4.06 | 30.25 | 2.20 | 0.148 |
| Species x shading | 2 | 0.01 | 0.01 | 3.85 | 0.10 | 0.903 |  | 7.41 | 3.71 | 5.70 | 2.01 | 0.218 |
| Direction x shading | 2 | **3.85** | **1.93** | **21.51** | **40.88** | **<0.001** |  | 4.25 | 2.13 | 29.86 | 1.16 | 0.329 |
| Species x direction x shading | 2 | 0.02 | 0.01 | 21.51 | 0.24 | 0.787 |  | 4.99 | 2.49 | 29.86 | 1.35 | 0.273 |

At the later developmental stage, the net flow of carbon in the full-light treatment was near zero in both species. However, the two species significantly differed in their response to shading treatment. While shading of daughters seemed to promote translocation toward daughters in both species, they responded differently to shading of mothers. In *F. viridis*, shading of mothers increased carbon translocation from daughters to mothers. In contrast, in *P. reptans*, shading of mothers reduced carbon translocation from daughters to mothers to undetectable values, while it did not markedly alter translocation to daughters (Table 1, Fig. 4, see Fig. S4 for individual values).

In relative terms, mother ramets of *F. viridis* and *P. reptans* in the full-light treatment exported on average 5.7 and 2.6% of assimilated carbon, respectively, to young daughters at the early developmental stage. At the later stage, the exported fractions were generally lower. Mother ramets of *F. viridis* and *P. reptans* exported 2.4% and 1.5% of assimilated carbon to shaded daughters, respectively. Daughters of *F. viridis* and *P. reptans* exported 4.7% and 0% of assimilated carbon to shaded mothers, respectively (Fig. 4). For detailed information on the exported proportions, see Table S2 in Supplementary Information.

### Nitrogen translocation

At the early developmental stage, nitrogen was only translocated toward daughters in both species, translocation from daughters to mothers was not detectable. *P. reptans* mothers translocated significantly more nitrogen to daughters than *F. viridis* mothers, in proportion to their higher nitrogen uptake. For both species, translocation to daughters was highest in the full-light treatment and lowest when the daughters were shaded (Table 1, Fig. 5).

**Figure 5.**
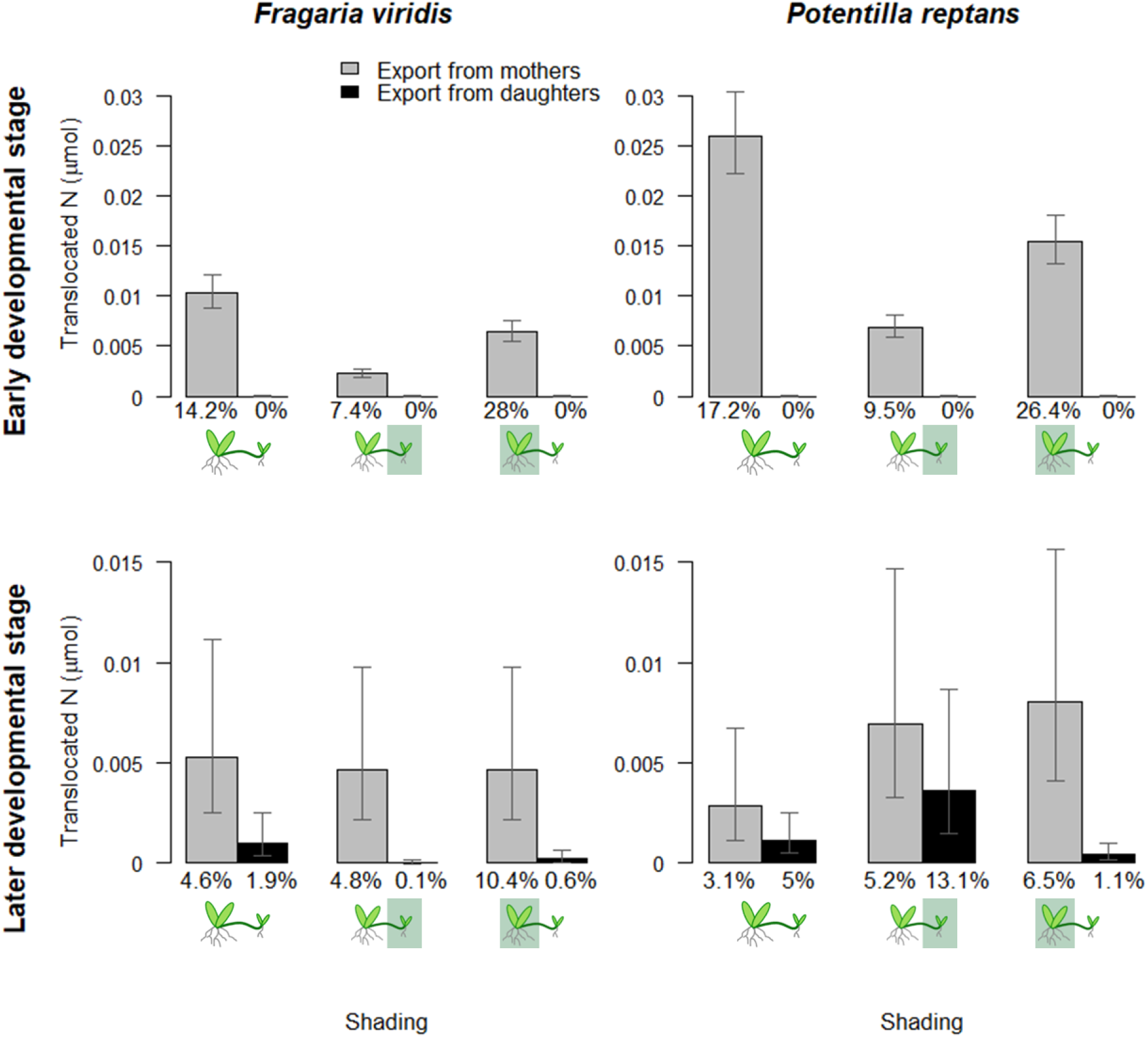
Absolute amounts of translocated ^15^N (means and SEM) from labelled mother to daughter ramets and from labelled daughter to mother ramets. Numbers under the bars indicate mean fractions of ^15^N exported from labelled ramets toward unlabelled ramets. Note that y-axis range for the later developmental stage is half of that for the early developmental stage. Values of absolute amounts are based on linear models with logarithmic transformation (values were back-transformed).

At the later developmental stage, nitrogen translocation toward daughters was still significantly higher in both species. Translocation to mothers was detectable, at least in some plants, and *P. reptans* tended to have higher translocation to mothers. However, differences between the species and shading treatments were not significant due to high variation (Table 1, Fig. 5).

The mean proportions of labelled nitrogen exported from mother ramets to early-stage daughters under homogeneous conditions were 14.2% for *F. viridis* and 17.2% for *P. reptans*. These proportions were modified by shading; with the highest proportions exported from shaded mothers and the lowest from unshaded mothers to shaded daughters. In the later stage, the mean exported proportions of nitrogen were 4.6 and 3.1% for mother ramets of *F. viridis* and *P. reptans,* respectively, under homogeneous conditions. These proportions increased to 10.4 and 6.5% when the mothers were shaded. The mean proportions exported by daughters ranged from 0.1% to 13.1% of labelled nitrogen, the latter exported by shaded *P. reptans* daughters (Fig. 5). See Table S2 in Supplementary Information for further details.

## Discussion

Using a pulse-chase labelling experiment, we showed transition from an early to a later pattern of resource sharing between mother and daughter ramets and identified two different later-stage patterns of carbon translocation in the two relative clonal species growing in habitats with different ecological regimes. Specifically, in *F. viridis,* carbon was translocated to shaded ramets at the later developmental stage. Therefore, the behaviour of *F. viridis* was consistent with the Equalisation strategy, as we expected. In contrast, *P. reptans* daughters did not export any carbon to shaded mothers and the translocation between *P. reptans* mothers and shaded daughters was less strongly daughter-directed than in *F. viridis*. The translocation pattern in *P. reptans* was therefore consistent with the expected Acropetal translocation strategy. A very similar pattern of carbon sharing was found in our previous experiment with *A. stolonifera* (Fig. 6). Nitrogen translocation was mainly acropetal and its pattern did not differ significantly between *F. viridis* and *P. reptans*, as discussed below.

**Figure 6.**
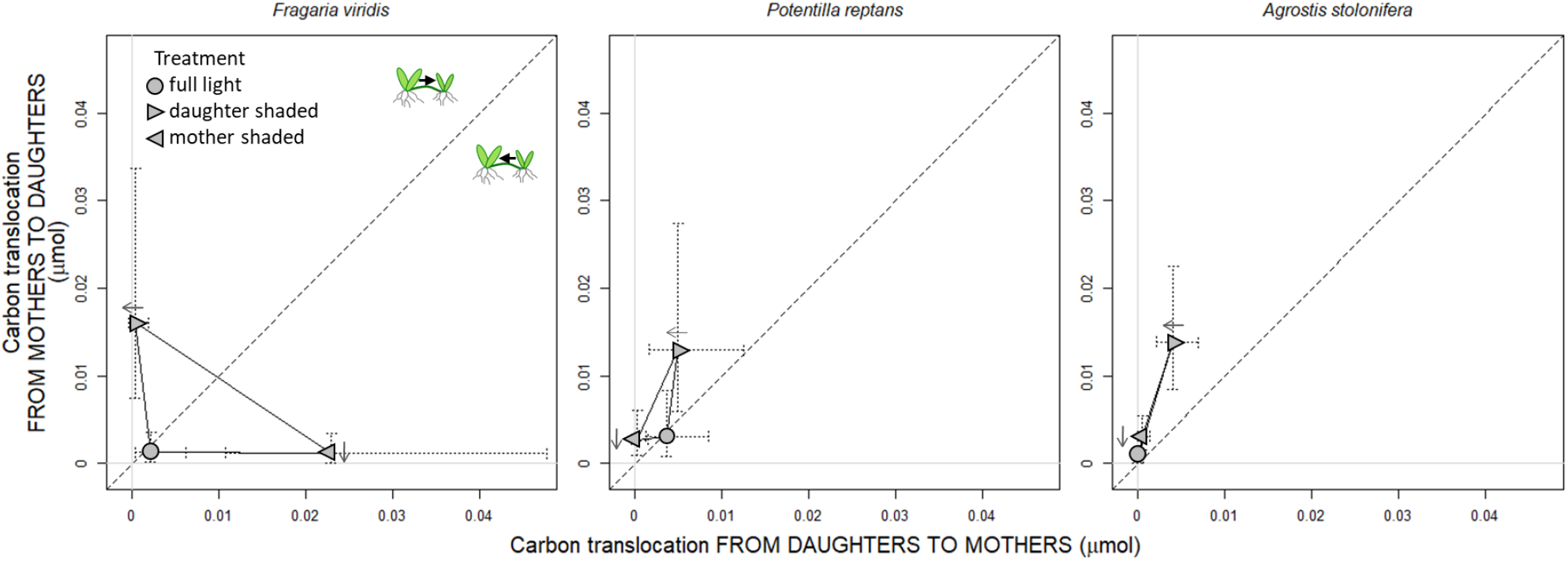
Two-way carbon translocation plots for the three examined species at the later developmental stage of daughter ramets (Duchoslavová & Jansa, 2018). F. viridis is a species of nutrient-poor grasslands, whereas P. reptans and A. stolonifera can grow in more nutrient-rich habitats with a higher degree of light competition. Means and SEM are depicted. Arrows next to the points indicate which values have possibly been overestimated by the labelling method, and the direction where more realistic values may lie.

### Early developmental stage of ramets

Physiological integration between mother and daughter ramets is inevitable at the early developmental stage, especially for soil-borne resources, as daughters’ resource-acquiring organs are still developing (e.g., Hartnett & Bazzaz, 1983; Duchoslavová & Jansa, 2018; Ma *et al*., 2021). In our experiment, shading of one ramet had very similar effect on biomass of both ramets suggesting high interdependence of ramets at the early developmental stage. Daughter ramets of both species studied were markedly smaller than mothers, had a lower uptake capacity for carbon and a very low (but detectable) uptake capacity for nitrogen. Accordingly, carbon and nitrogen translocation was predominantly directed to daughters in both species. However, shaded mothers did not appear to be a relatively stronger source of carbon than unshaded daughters even at the early developmental stage, as the translocation to the shaded mothers increased markedly, resulting in zero net carbon translocation. The acropetal nitrogen translocation followed the expected source-sink relations, with the lowest absolute translocation toward slower growing shaded daughters. Highest fractions of assimilated nitrogen were exported from shaded mothers to unshaded daughters, presumably to support their photosynthetic capacity (Saitoh et al. 2006), although the absolute amounts did not exceed translocation in the homogeneous treatment. Contrary to the carbon translocation, nitrogen translocation was determined by nitrogen uptake and relative size of ramets. In summary, shading conditions had significant effects on the translocation of both carbon and nitrogen during early development, which has rarely been investigated (Duchoslavová & Jansa, 2018).

### Later developmental stage of ramets

At the later developmental stage of ramets, daughter ramets were still significantly smaller than mothers, but generally had comparable carbon uptake capacity. Our results showed zero net translocation of carbon in homogeneous conditions, as did the previous study on *A. stolonifera* (Duchoslavová & Jansa, 2018). Interestingly, low amounts of carbon were still exchanged in both directions in some plants under homogeneous conditions, which was observed before in *Eichhornia crassipes* (Alpert *et al*., 1991). Persistent directed translocation of carbon occurred only under heterogeneous conditions, consistent with previous findings (Saitoh *et al*., 2002; Wang *et al*., 2021). Our results further suggest that bidirectional resource equalisation is only one of multiple possible translocation strategies in heterogeneous conditions. *F. viridis* translocated carbon toward shaded ramets regardless of their developmental position, in accordance with the Equalisation strategy. In contrast, carbon translocation toward shaded mother ramets stopped completely in *P. reptans* or *A. stolonifera* (Duchoslavová & Jansa, 2018). When only daughters were shaded, the net translocation flow was slightly directed toward daughters in these species (Fig. 6). Thus, these two species did not support developmentally older ramets and they seem to support developmentally younger ramets growing in less favourable conditions less extensively than *F. viridis*. This pattern is in accordance with the proposed Acropetal strategy of late resource sharing. The lack of support of older, resource-limited ramets in some species has previously been suggested by growth experiments (e.g., Xiao *et al*., 2011), but not demonstrated by labelling studies.

Regarding carbon source-sink relationships, larger and unshaded ramets presumably formed stronger sources whereas growing tissues formed main sinks for carbon (and nitrogen). Both the mother and daughter ramets were composed of rooted rosettes of leaves and unrooted stolons with leaves. As the rosettes presumably did not grow extensively at the later developmental stage, the unrooted stolons may have exhibited the most intensive growth. However, the role of the unrooted stolons in carbon source-sink relationships is speculative as we did not measure their carbon import and export separately. While the unrooted stolons may have acted as strong sinks driving the translocation (Ginzo & Lovell, 1973, Alpert 1999, Golovko et al. 2004), they may have been photosynthetically self-sufficient due to their developed leaves. Accordingly, several cases of unrooted and poorly rooted daughters tended to show higher carbon translocation to mothers (Fig. S4). This suggests an independence of unrooted stolons in carbon uptake, consistent with our previous findings in *Agrostis stolonifera* (Duchoslavová & Jansa, 2018).

Differences in nitrogen uptake between mothers and daughters at the later developmental stage remained more pronounced than in the case of carbon uptake, especially when daughters were shaded, and were driven by root mass of labelled ramets. Shading thus reduced nitrogen availability for ramets via its effect on growth (see also Freschet et al., 2018), although nitrogen levels were not manipulated in the experiment. The lower uptake of nitrogen in daughters, together with strong nitrogen sinks in the growing tissues of daughters, likely led to prevailing nitrogen translocation toward daughters in both species. Nitrogen translocation showed high variability, did not significantly differ between the two species and, in contrast to the early developmental stage, was not significantly affected by shading at the later developmental stage. This result did not confirm our expectations nor an observed effect of light heterogeneity on nitrogen translocation in *Sasa palmata*, where enhanced nitrogen translocation toward ramets in open patches likely increased the photosynthetic activity of illuminated leaves (Saitoh *et al*., 2006).

### Ecological consequences of the observed translocation strategies

The late-stage carbon sharing strategies proposed in this study are manifested under horizontal light heterogeneity, which may occur in patchy and sparse vegetation or in gaps in vegetation created by local disturbances. The two observed strategies may reflect overall availability of belowground resources and the level of aboveground competition typically experienced by a species. Maintaining older ramets with developed root systems through the Equalization translocation strategy may help plants to cope with limitation by belowground resources. In addition, under lower aboveground competition, it may be advantageous to support shaded ramets, which have a higher chance of overgrowing their neighbours. Support of younger shaded ramets may increase the competitive ability of clonal plants growing from an open area into the shaded conditions of a plant community, as was demonstrated in an experiment with *Fragaria chiloensis* (Wang *et al*., 2021). On the other hand, clonal growth is a way to colonise gaps in vegetation (Macek & Lepš, 2003; Kohler *et al*., 2006; Vítová *et al*., 2017), and the Acropetal translocation strategy may accelerate gap exploration and colonisation, especially when belowground resources are not limiting. We thus expect the Acropetal translocation strategy to be associated with a higher degree of light-foraging behaviour. Accordingly, *P. reptans* has been shown to adjust its growth response to the height and density of simulated neighbours, preferring lateral spread in the presence of tall neighbours (Gruntman *et al*., 2017).

However, the proposed connection between translocation strategy and habitat conditions, which implies different selection pressures, needs further testing by a comparative study. Although closely related, the two species also differ in other aspects, such as the positioning of flowers on stolons (*P. reptans*) or on rooted rosettes of leaves (*F. viridis*). This could lead to an alternative explanation of the different carbon translocation patterns at the later developmental stage, with *F. viridis* supporting older shaded ramet to promote their sexual reproduction (Alpert 2002).

Enhanced translocation of carbon to shaded daughters at the later developmental stage could be an active strategy to cope with heterogeneity in light availability or a pattern caused by slower growth and the resulting different ontogenetic stage of the shaded ramets (Huber et al., 1999). However, it is not possible to distinguish between the effects of daughter size and shading as they strongly covary.

### Effect of translocation patterns on growth

Most clonal translocation studies have measured the growth characteristics of plants instead of using labelled-element translocation, which is more technically and financially demanding. The link between resource-sharing pattern and growth response is crucial for understanding the impacts of translocation strategies. Unfortunately, only a few studies have provided both tracing and growth data for the same experimental setup (see D’Hertefeldt & Jonsdottir, 1994; Xu *et al*., 2010, 2012; Duchoslavová & Jansa, 2018). Xu and collaborators (2010) obtained matching results in two creeping species for the integration effect on daughter-ramet growth and late-stage carbon translocation toward daughters. However, determining the translocation strategy from growth data is complicated in later developmental stages by initial acropetal translocation, which may overlap the effect of the later translocation pattern (Duchoslavová & Jansa, 2018). Thus, an experimental design with repeated measurements of growth that allow the estimation of the growth rate over the experimental period is needed to match observed translocation patterns with ramets’ actual growth (Abrahamson *et al*., 1991, Ma *et al*., 2021).

Nevertheless, estimating the impact of translocation on plant growth is important despite the lack of growth data. It may be difficult to interpret absolute amounts of translocated resources assimilated in short labelling pulses in terms of their impact on ramet growth and functioning. The proportions of recently assimilated resources that are exported may give a more relevant picture of the translocation impact. Pitelka and Ashmun (1985) consider an export of 1–5% of a resource to have a significant effect on growth, reproduction, or survival. In conditions distinguishing the two strategies in our experiment (i.e. in the later developmental stage under heterogeneous light), *Fragaria* translocated on average 2.4 and 4.7% of assimilated carbon to shaded daughter and mother ramets, respectively. This was comparable to its initial maternal export constituting 4.9 to 6.5% of assimilated carbon. We thus consider this later translocation in *Fragaria* to have a significant effect on plant functioning. However, later-stage translocation flows in *Potentilla* were less clear – carbon translocation between mothers and shaded daughters occurred in both directions and only 0.5% of assimilated carbon was translocated from shaded mothers. Effect of such translocation rates on growth of established daughters may be questionable and, therefore, the most distinctive characteristic of *Potentilla* resource-sharing pattern is the absence of translocation toward shaded developmentally older ramets.

### Methodological issues

Despite the number of translocation studies, it is difficult to identify translocation strategies in the literature. We see two major reasons for this. First, translocation in the Equalisation or Acropetal translocation strategy does not differ in homogeneous conditions. Therefore, it is not possible to distinguish these two translocation strategies in studies that do not involve environmental gradients. Nevertheless, many tracing studies use homogeneous conditions that do not allow the identification of resource-sharing strategies (e.g., Ginzo & Lovell, 1973; Alpert, 1996; D’Hertefeldt & Jonsdottir, 1999). Further, our results demonstrate the necessity of bidirectional tracing for the recognition of translocation strategies. Counterintuitively, basipetal translocation of carbon showed higher plasticity than acropetal translocation in the studied species; acropetal carbon translocation alone (Fig. 4, grey bars) would not reveal marked differences. However, many studies have traced labelled resources in only one direction, and important parts of the story may thus remain hidden (Saitoh *et al*., 2006; Xu *et al*., 2010). Similarly, the necessity of bidirectional tracing for estimating the carbon budget was recognised in a system of plant and mycorrhizal fungi (Cameron *et al*., 2008).

Bidirectional tracing of carbon translocation between ramets in different light regimes, however, poses some methodological challenges. Photosynthesis is generally slower in shaded conditions, resulting in lower carbon uptake by shaded ramets. Labelling under shade may be logistically demanding and concentration of labelled carbon in plant tissues may be too low to detect translocation flow. Therefore, we decided to label the plants in full-light conditions although the uptake of carbon by shaded ramets was thus likely overestimated to some extent. Consequently, the absolute values of carbon translocation from the shaded ramets may also have been overestimated, although there was no significant relationship between carbon uptake and export (R^2^ = 0.04). The possible overestimation is reflected in the result interpretation, and it could not have affected observed differences between the species. Moreover, the relative carbon translocation values, which were unlikely to be affected by two hours of labelling, merely confirmed the observed patterns (Fig. 4).

To enhance the informative power of the results, we handled the background values of ^13^C and ^15^N concentrations in an innovative manner. Instead of using the mean concentration values of reference plants, we used detectable enrichment threshold value to filter out the background concentrations (see Methods for further details). Values below this threshold were considered zero. This approach resulted in less statistically significant but more credible results.

Although we traced the translocation of both carbon and nitrogen in our experiment, we only manipulated the availability of light. It is likely that translocation strategies under environmental heterogeneity in soil-borne nutrients also vary between species and may be influenced by the productivity of the species’ habitats. However, competition for nutrients does not switch from symmetric to asymmetric, as in the case of light competition. Translocation strategies under nutrient heterogeneity may differ from those found under light heterogeneity due to the different nature of competition for these resources.

## Conclusion

Our experiment demonstrated two distinct carbon sharing strategies between established ramets in two closely related stoloniferous species under heterogeneous light conditions. Furthermore, we showed that the light gradient affects the translocation of both carbon and nitrogen at the early developmental stage. We articulated several hypotheses regarding the relationship between translocation strategies and the species’ typical environment. Comparative studies involving multiple species are necessary to test these aspects. We see the experiment presented here as a first step towards this goal. It is important to note that accurate comparisons can only be made through bidirectional translocation measurements, which involves measuring translocation from both mother to daughter and daughter to mother.

## Acknowledgements

We are grateful to Tomáš Herben, Peter Alpert and two anonymous reviewers for their helpful comments.

## Author contributions

Jana Duchoslavová planned the experimental design, conducted the experiment, analysed the data and wrote the manuscript. Jan Jansa advised on the experimental design and provided the elemental and isotopic analyses. Both authors contributed critically to the drafts and gave final approval for publication.

## Data availability

The data that support the findings of this study are openly available in Dryad.

