## Supplementary Data for "Strategies of resource sharing in clonal plants: A conceptual model and an example of contrasting strategies in two closely related species"

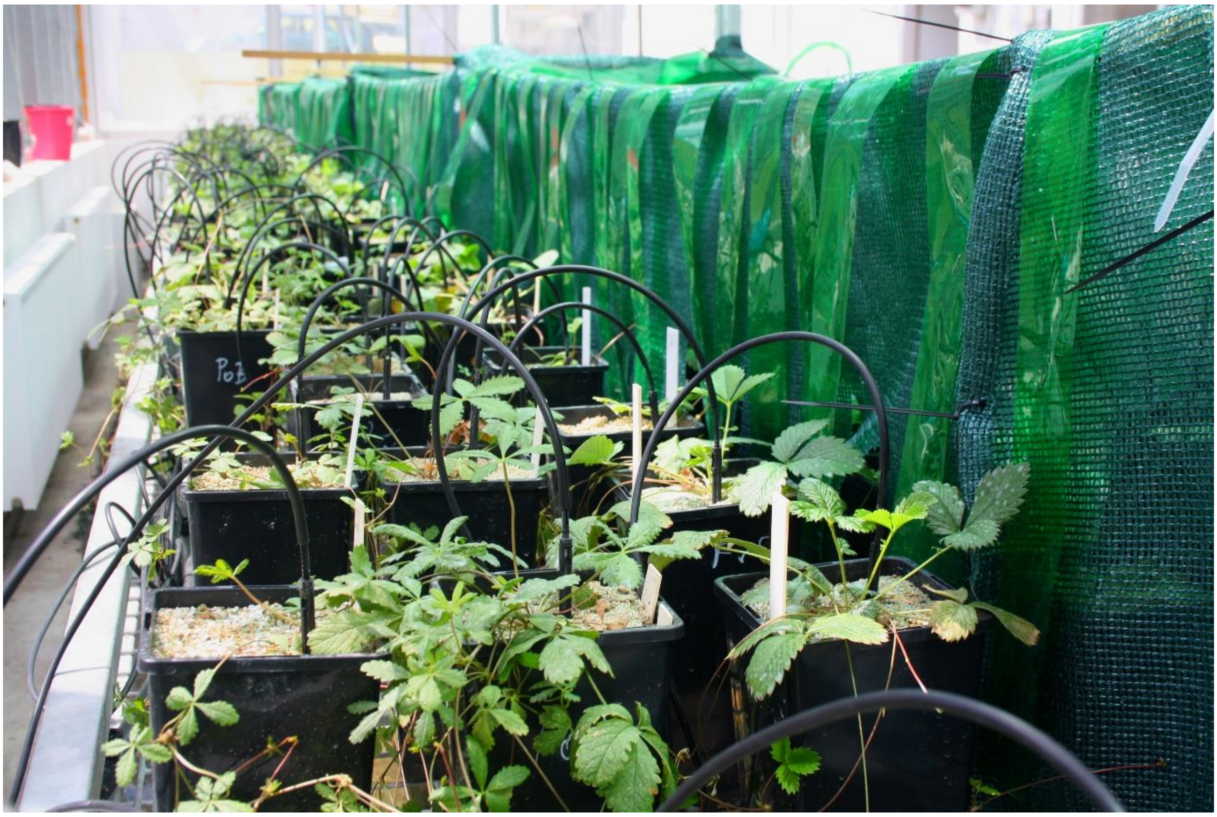

Figure S1. *Photograph of experimental setup. Full-light treatment is on the left; treatments with mother or daughter shaded are on the right, with one ramet under the shading cloth.*

Table S1. *ANOVA table of the linear mixed-effects models of ramet biomass. Significant effects are depicted in bold.*

**Biomass of ramets**

| | | **First harvest** | | | | **Second harvest** | | | |
|---|---|---|---|---|---|---|---|---|---|
| | d.f. | SumSq | DenDF | Fvalue | Pr(>F) | SumSq | DenDF | Fvalue | Pr(>F) |
| Initial number of leaves | **1** | **0.69** | **78.52** | **9.04** | **0.004** | 0.21 | 84.84 | 1.60 | 0.210 |
| Initial number of stolons | 1 | 0.17 | 74.00 | 2.15 | 0.147 | 0.08 | 84.33 | 0.60 | 0.441 |
| Ramet | **1** | **47.82** | **75.48** | **622.53** | **<0.001** | **8.34** | **81.27** | **62.95** | **<0.001** |
| Shading | **2** | **4.00** | **78.52** | **26.02** | **<0.001** | **3.35** | **82.64** | **12.64** | **<0.001** |
| Species | 1 | 0.12 | 5.53 | 1.58 | 0.259 | 0.09 | 5.26 | 0.67 | 0.450 |
| Ramet x shading | 2 | 0.33 | 75.48 | 2.13 | 0.126 | **2.40** | **81.27** | **9.07** | **<0.001** |
| Ramet x species | 1 | 0.08 | 75.48 | 0.99 | 0.323 | **0.65** | **81.27** | **4.92** | **0.029** |
| Shading x species | 2 | 0.14 | 78.52 | 0.92 | 0.402 | **1.18** | **83.18** | **4.44** | **0.015** |
| Ramet x shading x species | 2 | 0.13 | 75.48 | 0.82 | 0.442 | 0.10 | 81.27 | 0.38 | 0.687 |

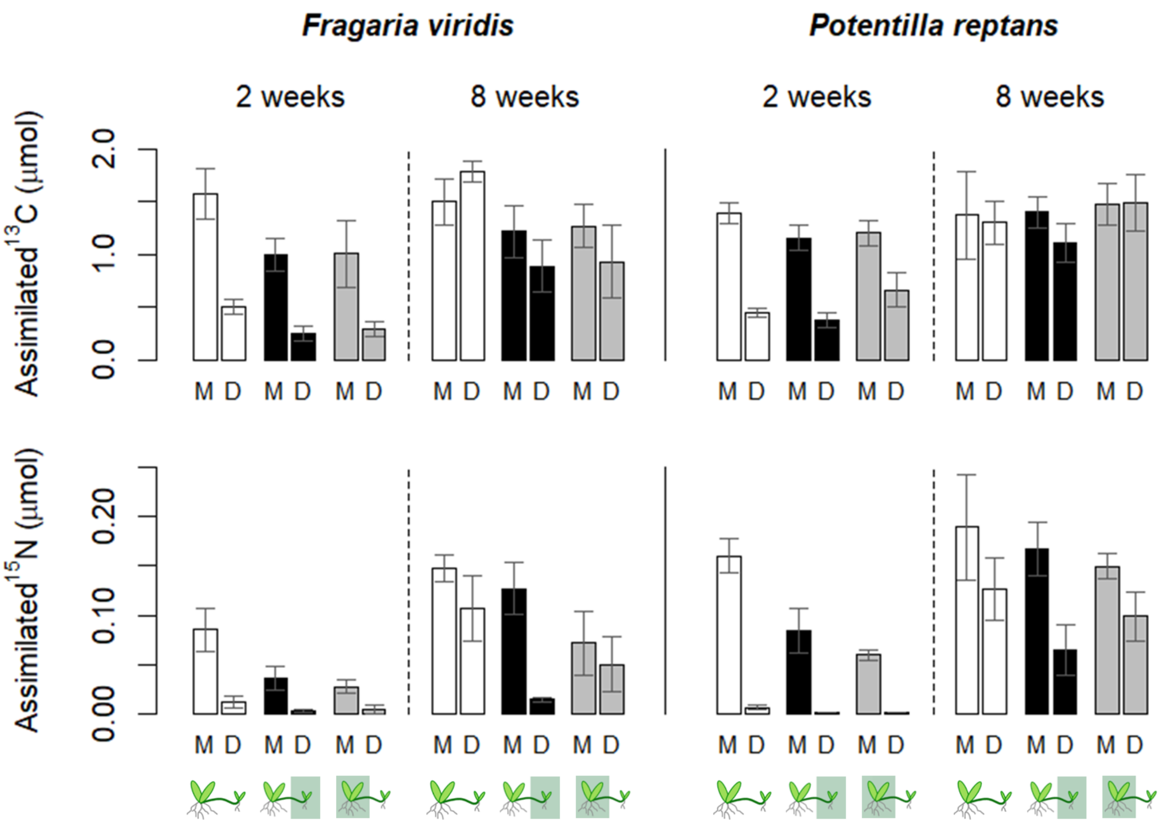

Figure S2. *Total assimilated* $^{13}C$ *and* $^{15}N$ *by labelled mother and daughter ramets at early and later developmental stages of daughter ramets (i.e. 2 and 8 weeks after daughter initialisation). Means and SEM are depicted.*

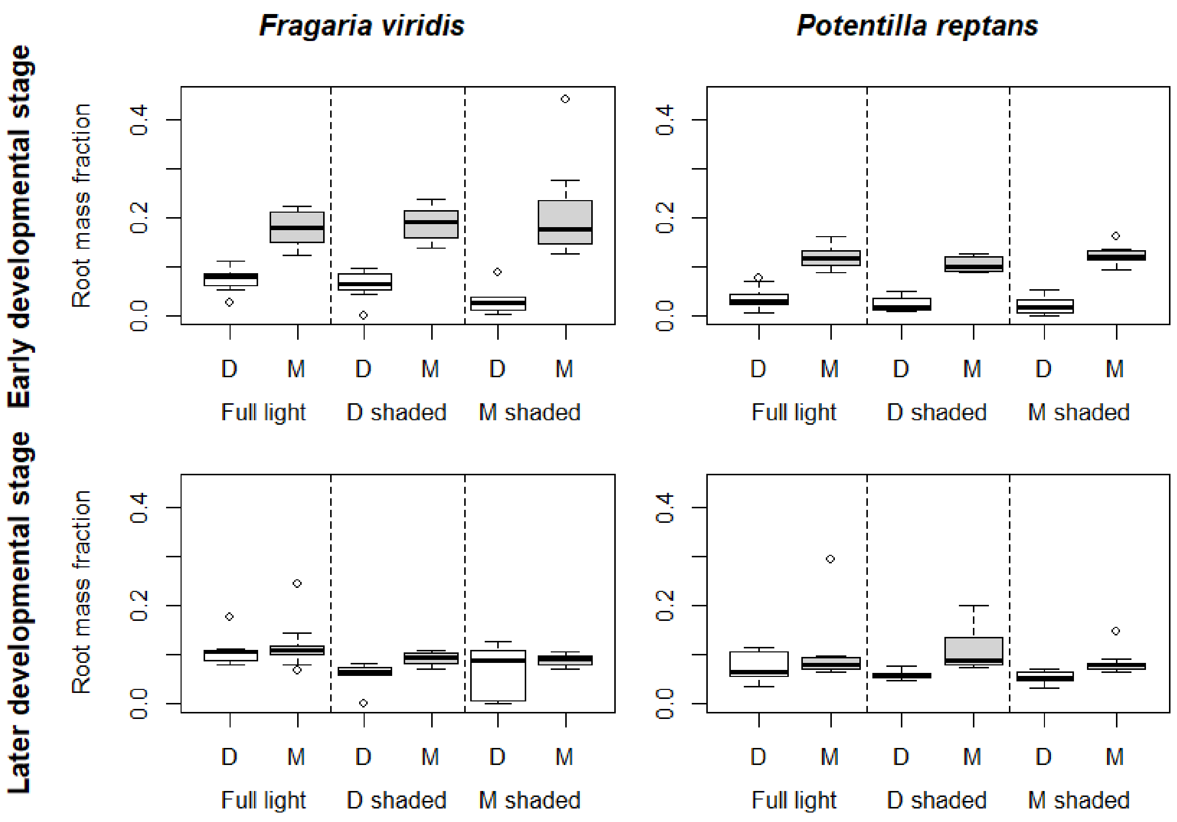

Figure S2. *Root mass fraction of ramets at the early and later developmental stages (2 and 8 weeks after daughter initialisation, respectively). White boxplots depict daughter ramets and grey boxplots depict mother ramets. The boxes represent medians and quartiles, the whiskers shows data range without outliers.*

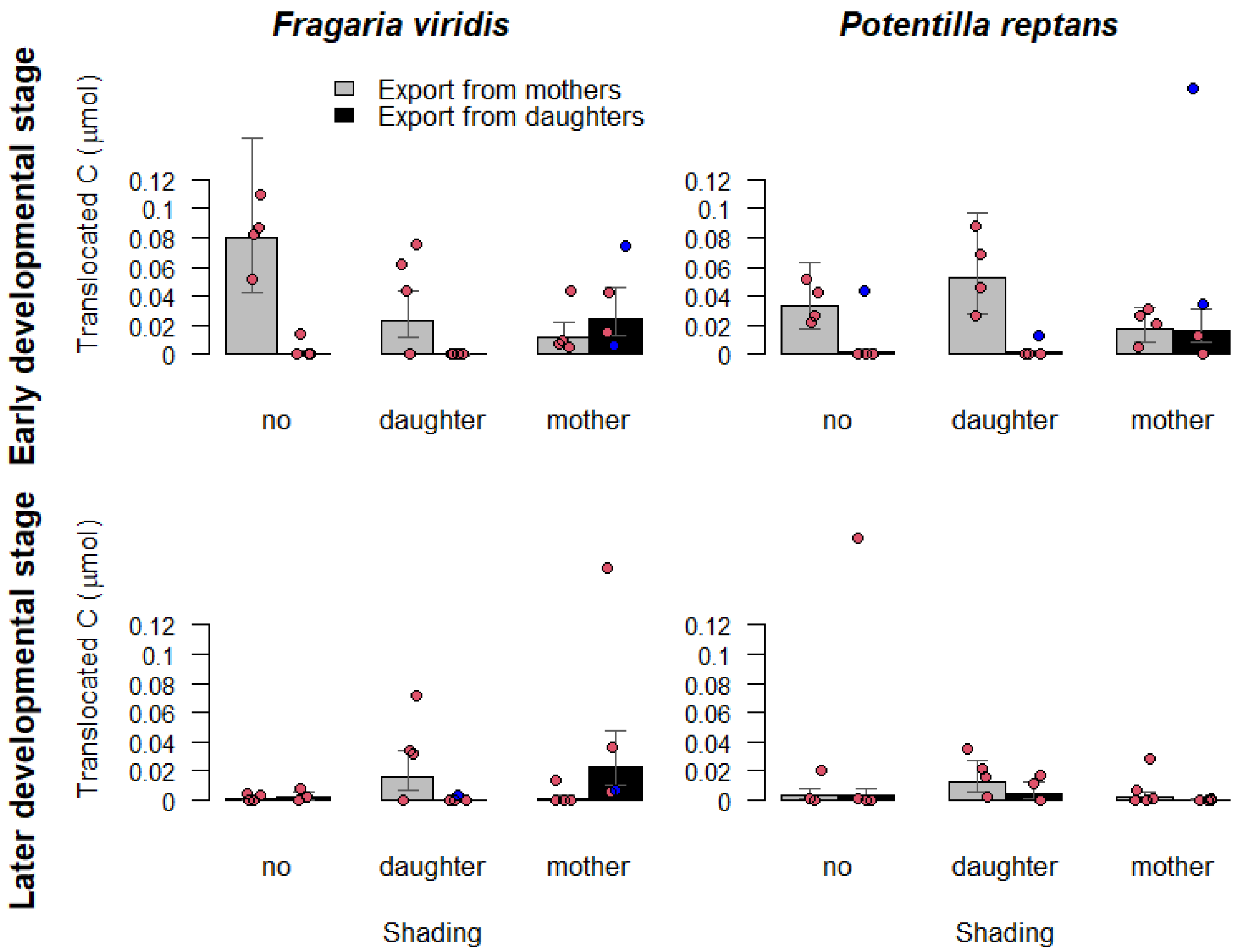

Figure S4. *Individual values of translocated $^{13}C$. Points represent individual values, while bar plots represent mean values and SEM based on log-transformed data. The blue points represent values of $^{13}C$ translocated from daughters with underdeveloped roots, as indicated by their low N uptake.*

Table S2. *Fractions of $^{13}C$ and $^{15}N$ exported from labelled ramets toward unlabelled ramets.*

| | *Fragaria viridis* | | | | *Potentilla reptans* | | | |
|---|---|---|---|---|---|---|---|---|
| **Exported carbon** | | | | | | | | |
| | **From mothers** | | **From daughters** | | **From mothers** | | **From daughters** | |
| **First harvest** | mean | SE | mean | SE | mean | SE | mean | SE |
| Full light | 5.73% | 1.20% | 0.60% | 0.59% | 2.62% | 0.52% | 2.05% | 2.05% |
| Daughter shaded | 4.91% | 1.74% | 0.00% | 0.00% | 5.33% | 1.47% | 0.62% | 0.58% |
| Mother shaded | 6.46% | 5.89% | 11.17% | 3.92% | 1.66% | 0.43% | 6.37% | 3.57% |
| **Second harvest** | mean | SE | mean | SE | mean | SE | mean | SE |
| Full light | 0.12% | 0.07% | 0.21% | 0.15% | 1.07% | 1.01% | 2.39% | 2.34% |
| Daughter shaded | 2.39% | 0.84% | 0.29% | 0.29% | 1.48% | 0.59% | 1.04% | 0.64% |
| Mother shaded | 0.29% | 0.26% | 4.68% | 1.81% | 0.49% | 0.35% | 0.02% | 0.02% |
| **Exported nitrogen** | | | | | | | | |
| | **From mothers** | | **From daughters** | | **From mothers** | | **From daughters** | |
| **First harvest** | mean | SE | mean | SE | mean | SE | mean | SE |
| Full light | 14.16% | 2.58% | 0.00% | 0.00% | 17.17% | 2.76% | 0.00% | 0.00% |
| Daughter shaded | 7.40% | 1.14% | 0.00% | 0.00% | 9.45% | 1.88% | 0.00% | 0.00% |
| Mother shaded | 28.01% | 7.44% | 0.02% | 0.02% | 26.38% | 2.49% | 0.00% | 0.00% |
| **Second harvest** | mean | SE | mean | SE | mean | SE | mean | SE |
| Full light | 4.60% | 1.40% | 1.88% | 1.31% | 3.09% | 2.00% | 5.00% | 3.78% |
| Daughter shaded | 4.80% | 1.97% | 0.09% | 0.09% | 5.21% | 1.83% | 13.14% | 7.43% |
| Mother shaded | 10.39% | 2.73% | 0.63% | 0.10% | 6.53% | 1.68% | 1.12% | 0.74% |
